## Supplementary information for "Dissecting crosstalk induced by cell-cell communication using single-cell transcriptomic data"

This file includes Supplementary Figures 1-7, Supplementary Tables 1-2, Supplementary Notes 1-4, and Supplementary References.

### Supplementary Figures

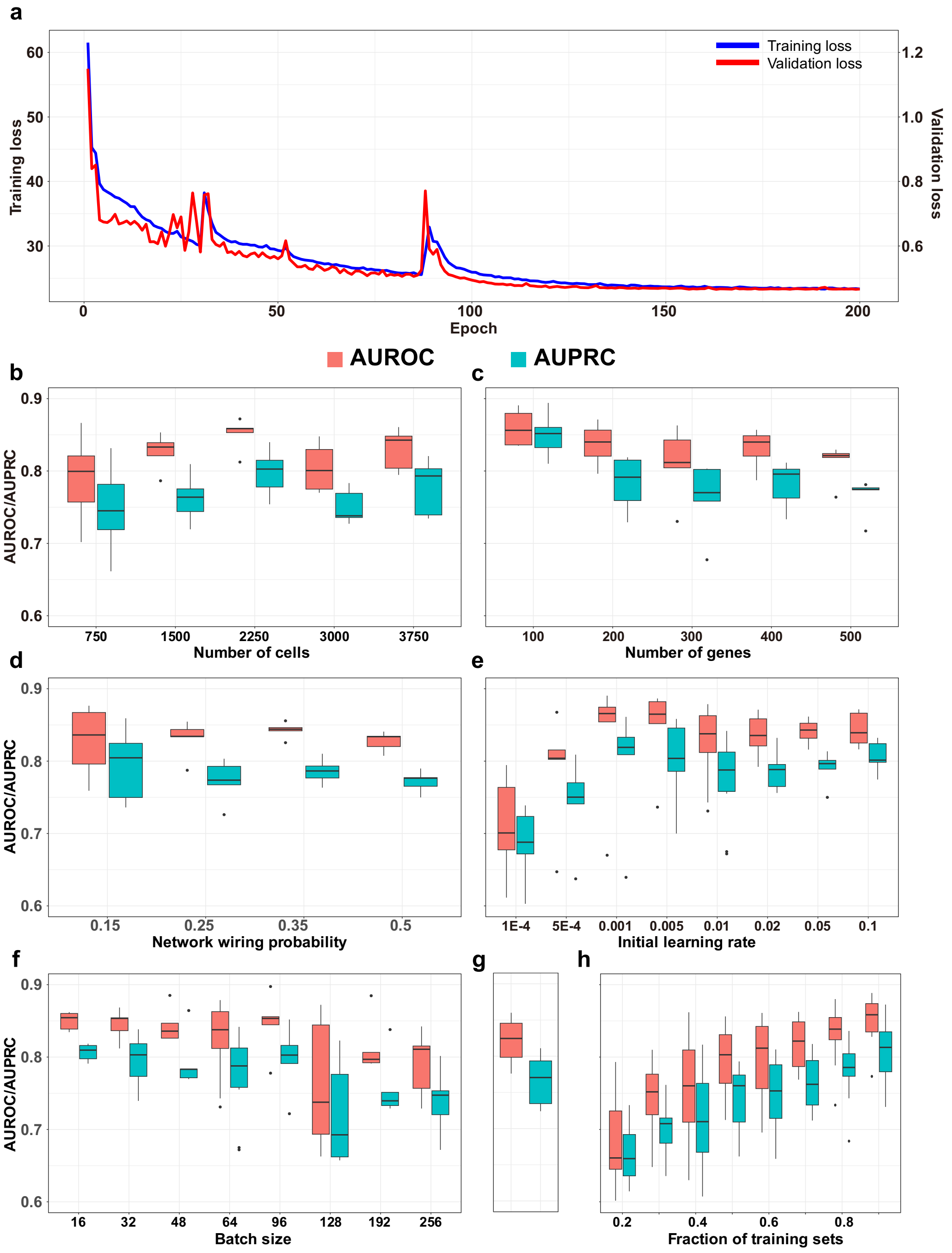


**Supplementary Figure 1**. The performance of SigXTalk on recovering the regulatory pathways under different conditions or hyperparameter settings. **a**. The curve of training and validation loss over epochs during a single training process. **b**. The effect of number of cells in the simulated dataset; **c**. The effect of number of genes in the simulated dataset; **d**. The effect of the density of the gene interaction network in the simulated dataset. The density is represented by the wiring probability of gene pairs; **e**. The effect of the initial learning rate on the performance; **f**. The effect of the initial learning rate on the performance; **g**. The effect of the choice of 10 random seeds on the performance; **h**. The effect of the training size used in the self-supervised training strategy on the performance, based on 10 independent test runs.

Boxplot elements: center line, median; box limits, upper and lower quartiles; whiskers, 1.5x interquartile range; points, outliers. Source data are provided as a Source Data file. All the tests on the effect of different conditions or hyperparameter settings are performed on 5 independent test runs unless specified.

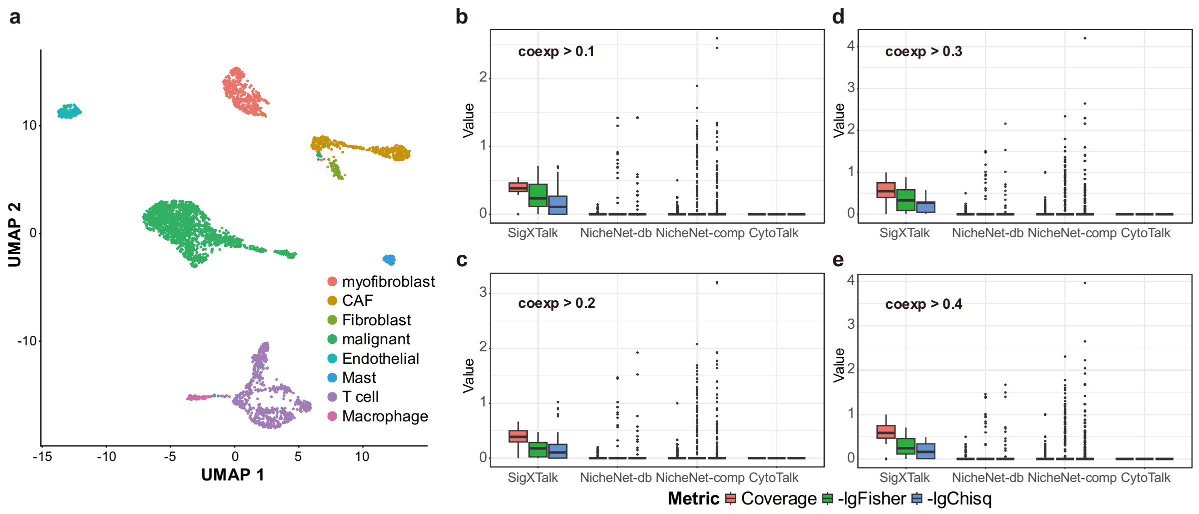


**Supplementary Figure 2**. Validation of SigXTalk using real dataset under different co-expression levels (coexp). **a**. Overview of the HNSCC dataset. Cells are visualized using Uniform Manifold Approximation and Projection (UMAP) as the dimensional reduction method; **b**-**e**. Performance of SigXTalk, NicheNet-db, NicheNet-comp and CytoTalk when the minimum co-expression level between signal and SSC is set to 0.1 (**b**, sample size: 42, 125, 382, 1 for the four methods), 0.2 (**c**, sample size: 42, 119, 366, 1 for the four methods), 0.3 (**d**, sample size: 42, 109, 337, 1 for the four methods), and 0.4 (**e**, sample size: 42, 83, 278, 1 for the four methods). Boxplot elements: center line, median; box limits, upper and lower quartiles; whiskers, 1.5x interquartile range; points, outliers. Source data are provided as a Source Data file.

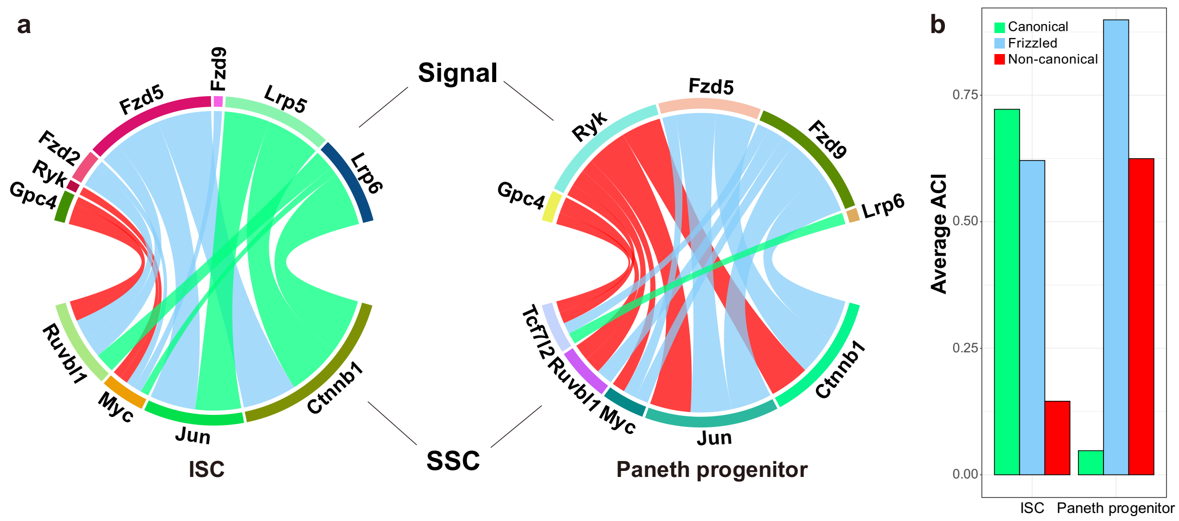


**Supplementary Figure 3**. The activation of pathways regulating Lgr5 calculated by individual-cell-based methods. **a**. The chord diagrams for the activation index (ACI) of each signal-SSC pair in intestinal stem cells (ISCs, left panel) and Paneth progenitors (right panel). The width of the edge is proportional to the average of ACI of all cells in the corresponding cell cluster; **b**. The bar charts for the average ACI values of each kind of Wnt signal (canonical, non-canonical, or frizzled signals that are shared by both canonical and non-canonical pathways) in ISCs and Paneth progenitors. Source data are provided as a Source Data file.

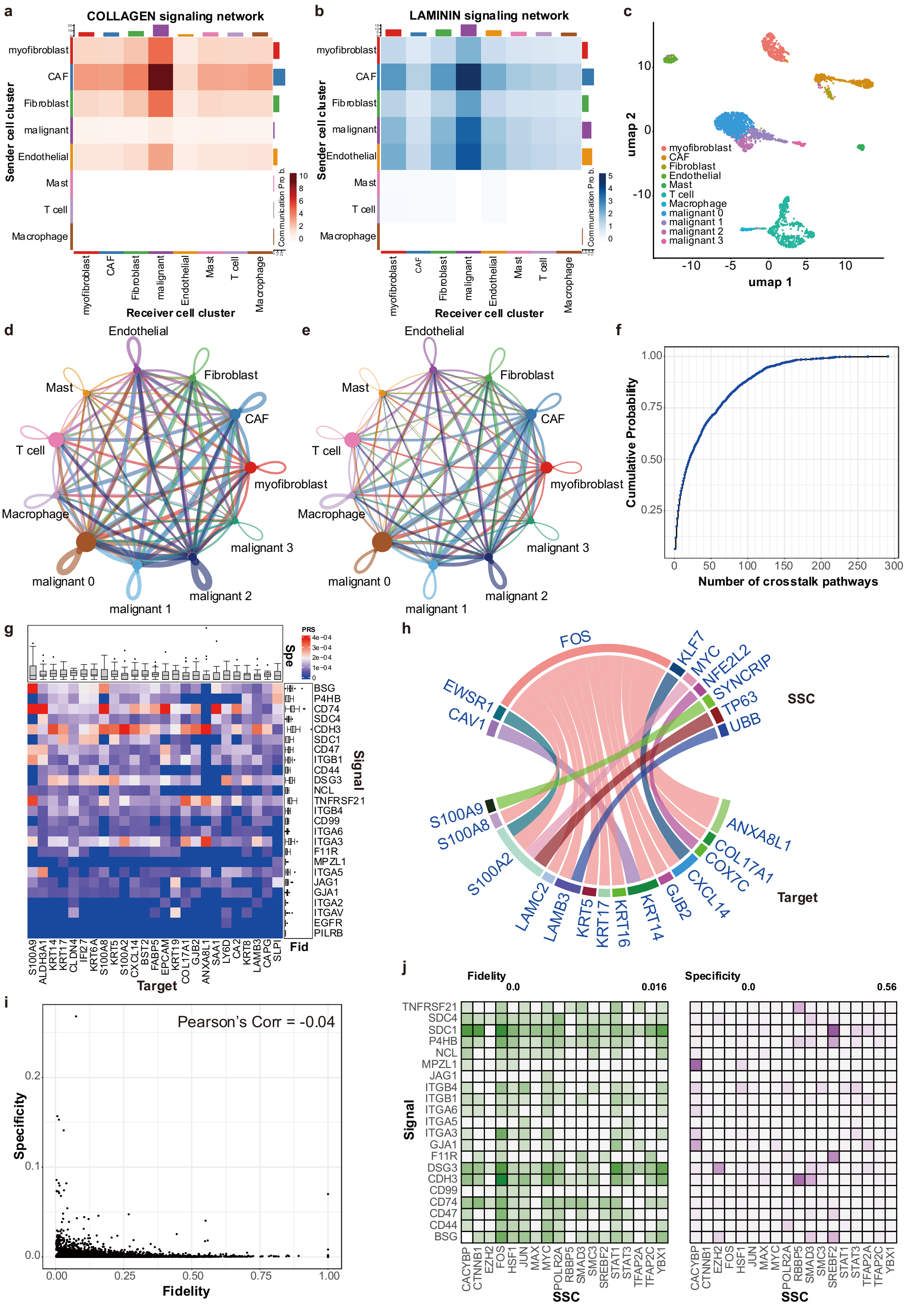


**Supplementary Figure 4**. Detailed crosstalk patterns in the HNSCC dataset. **a**. Heatmap of the Collagen-mediated cell-cell communication between cell clusters; **b**. Heatmap of the Laminin-mediated cell-cell communication between cell clusters; **c**. Overview of the HNSCC dataset after the malignant cells are subclustered. Cells are visualized using UMAP as the dimensional reduction method; **d-e**. The CCC patterns among cell clusters after subclustering the malignant. The width of each link stands for the number of Ligand-Receptor pairs (**d**) and communication strength (**e**) from the sender to the receiver; **f**. The cumulative density function of the number of crosstalk pathways for each target; **g**. Heatmap for the PRS value of each pathway that is mediated by *FOS*. The upper and right bar charts annotate the specificity (Spe) and fidelity (Fid) of the corresponding pathways, respectively (sample size: 25 for each target or signal); **h**. The chord diagram for the specificity values of the pathways regulated by the signal *CDH3*. The width of the edge is proportional to the specificity value. **i**. The scatter plot showing the relationship between the fidelity and specificity values of all the signal-target pairs. **j**. The heatmaps of the fidelity/specificity values of each pathway that regulates *KRT17*. Here the crosstalk module used for calculation of fidelity/specificity is defined by the pathways that share SSC+Target or Signal+SSC, respectively. Boxplot elements: center line, median; box limits, upper and lower quartiles; whiskers, 1.5x interquartile range; points, outliers. Source data are provided as a Source Data file.

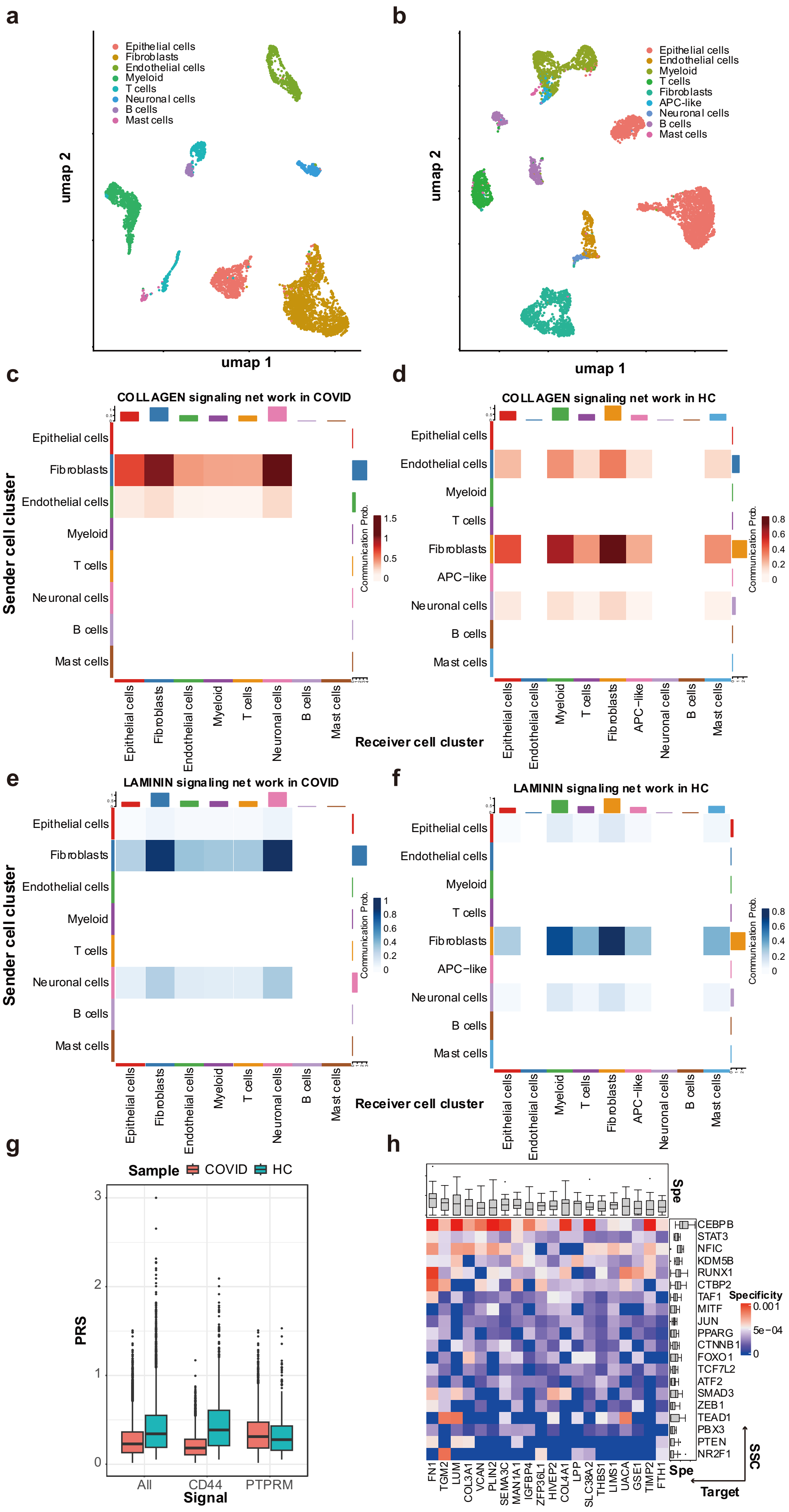
a
b
c
d
e
f
g
h
umap 1
umap 2
Epithelial cells
Fibroblasts
Endothelial cells
Myeloid
T cells
Neuronal cells
B cells
Mast cells
APC-like
COLLAGEN signaling net work in COVID
COLLAGEN signaling net work in HC
LAMININ signaling net work in COVID
LAMININ signaling net work in HC
Sender cell cluster
Receiver cell cluster
Communication Prob.
Sample
COVID
HC
PRS
Signal
All
CD44
PTPRM
Spe
SSC
Target
Specificity
CEBPB
STAT3
NFIC
KDM5B
RUNX1
CTBP2
TAF1
MITF
JUN
PPARG
CTNNB1
FOXO1
TCF7L2
ATF2
SMAD3
ZEB1
TEAD1
PBX3
PTEN
NR2F1
FN1
TGM2
LUM
COL3A1
VCAN
PLIN2
SEMA3C
MAN1A1
IGFBP4
ZFP36L1
HIVEP2
COL4A1
LPP
SLC38A2
THBS1
LIMS1
UACA
GSE1
TIMP2
FTH1

**Supplementary Figure 5**. The CCC and CCC-induced crosstalk patterns in the COVID dataset. **a**-**b**. Overview of the COVID dataset. Cells of the COVID (**a**) and the HC (**b**) sample are visualized using UMAP as the dimensional reduction method; **c-d**. Heatmap of the Collagen-mediated cell-cell communication between cell clusters in the COVID (**c**) and the HC sample (**d**); **e-f**. Heatmap of the Laminin-mediated cell-cell communication between cell clusters in the COVID (**e**) and the HC sample (**f**); **g**. The box diagram for the PRS values regulated by different signals in the COVID and HC sample. Sample size: All signals, 12,680 for COVID and 4,290 for HC; CD44, 4,154 for COVID and 1,140 for HC; PTPRM, 4,346 for COVID and 969 for HC. **h**. Heatmap of the PRS value of each pathway that is regulated by *ITGB1* in the COVID sample. The upper and right boxplots annotate the specificity (Spe) of the corresponding pathways. Sample size: 20 for each Target or SSC. Boxplot elements: center line, median; box limits, upper and lower quartiles; whiskers, 1.5x interquartile range; points, outliers. Source data are provided as a Source Data file.

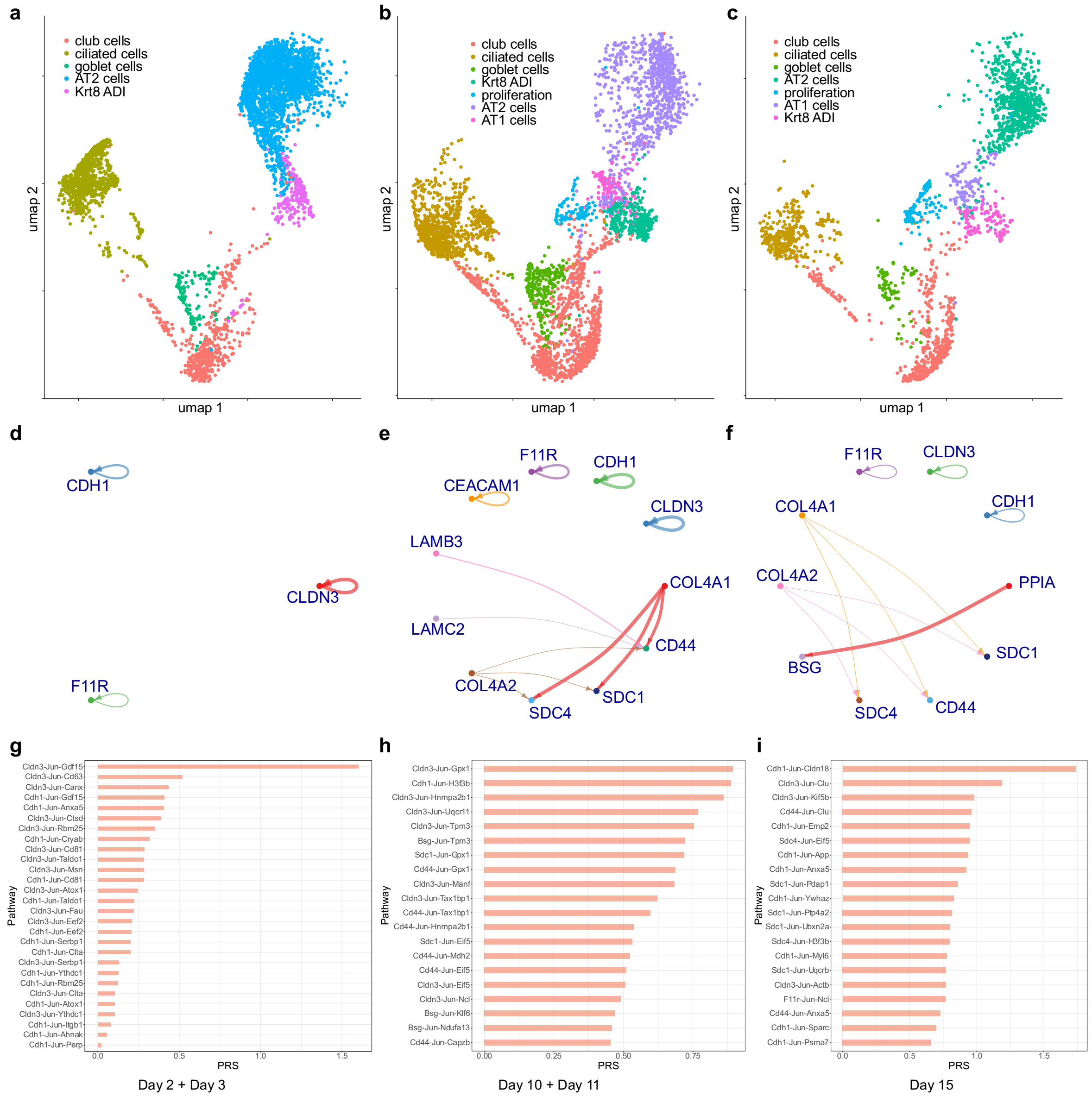

**Supplementary Figure 6**. Evolution of specific crosstalk pathways in the mouse lung dataset. **a-c**. Overview of the mouse lung dataset at Day 2+Day 3 (**a**), Day 10+Day 11 (**b**) and Day 15 (**c**). Cells of the three time points are visualized using UMAP as the dimensional reduction method; **d-f**. Circle plot of the interaction strength of CCC when the Krt8 ADI is set as the receiver cell cluster at Day 2+Day 3 (**d**), Day 10+Day 11 (**e**) and Day 15 (**f**). **g-i**. The bar charts for the pathway regulatory strength (PRS) values of the regulatory pathways that are mediated by *Jun*. Source data are provided as a Source Data file.

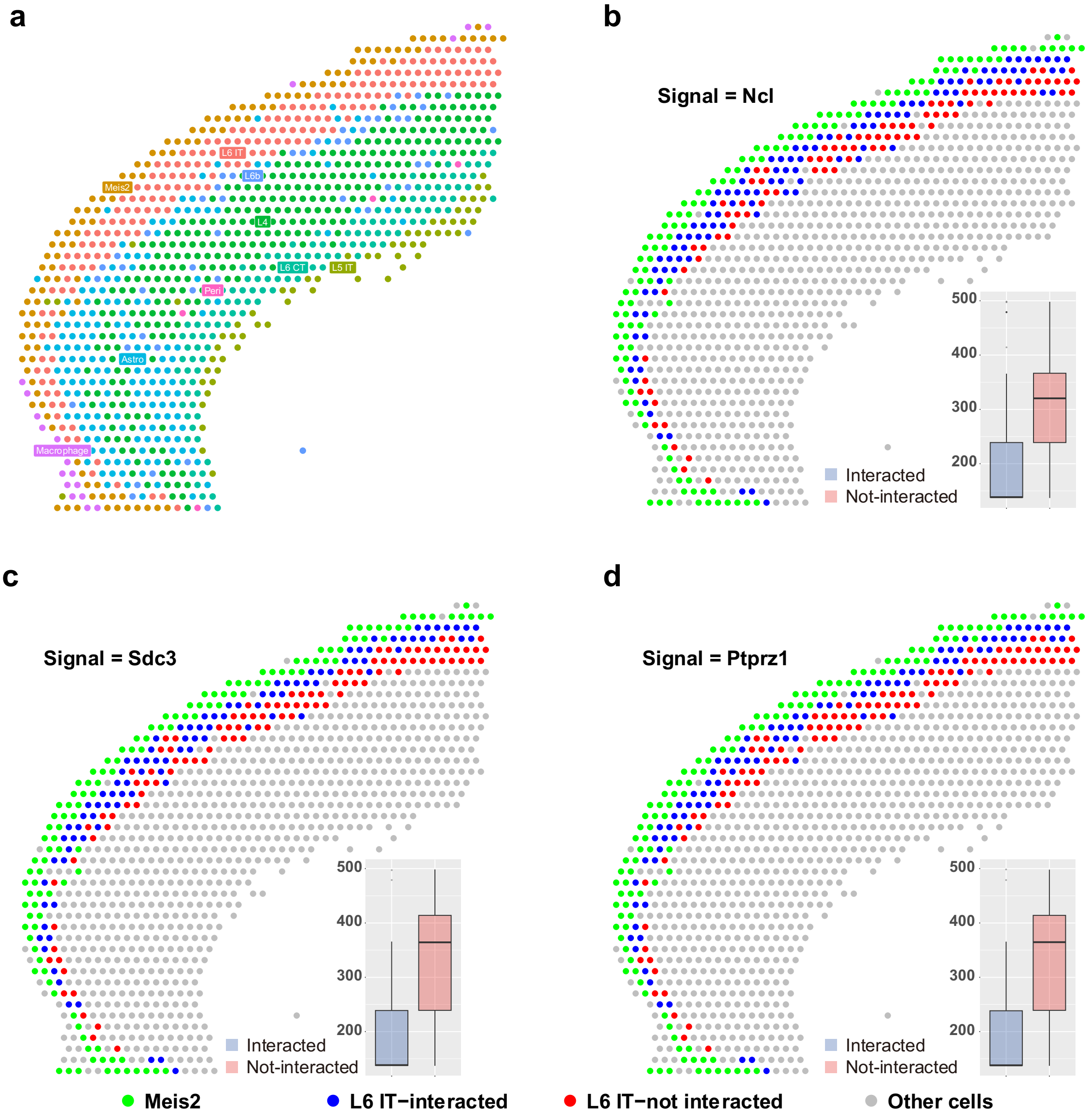


**Supplementary Figure 7**. Sub-clustering of the L6 IT cells in the mouse brain 10x Visium dataset induced by other signals. **a**. The spatial distribution of the cells of the dataset; **b**. The spatial plot of the L6 IT cells subclustered by the interactions with Meis2 cells, using the highly specific genes regulated by *Ncl* (sample size: 108 interacted and 88 not-interacted spots); **c**. The spatial plot of the L6 IT cells subclustered by the interactions with Meis2 cells, using the highly specific genes regulated by *Sdc3* (sample size: 98 interacted and 98 not-interacted spots). **d**. The spatial plot of the L6 IT cells subclustered by the interactions with Meis2 cells, using the highly specific genes regulated by *Ptprz1* (sample size: 94 interacted and 102 not-interacted spots). All the boxplots in the inner panels measure the distance from L6 IT cells to the Meis2 cells using the signal-induced subclustering results. Boxplot elements: center line, median; box limits, upper and lower quartiles; whiskers, 1.5x interquartile range; points, outliers. Source data are provided as a Source Data file.

### Supplementary Tables

**Supplementary Table 1.** The coverage of the predicted shared TFs out of all the ground truth shared TFs in different cell types

| **Cell type** | **Average coverage of methods** | | | | **P-value of two-sided t-test** | |
|---|---|---|---|---|---|---|
| | SigXTalk | NicheNet-db | NicheNet-comp | CytoTalk | SigXTalk vs. NicheNet-db | SigXTalk vs. NicheNet-comp |
| CAF | 0.367 | 0.011 | 0.035 | 0 | $1.3 \times 10^{-15}$ | $7.0 \times 10^{-15}$ |
| Malignant | 0.362 | 0.041 | 0.109 | 0 | $< 2.2 \times 10^{-16}$ | $< 2.2 \times 10^{-16}$ |
| Myofibroblast | 0.611 | 0.100 | 0.058 | 0 | $4.4 \times 10^{-3}$ | $< 2.2 \times 10^{-16}$ |
| T cell | 0.402 | 0.083 | 0.136 | 0 | $5.2 \times 10^{-4}$ | $2.8 \times 10^{-5}$ |
| Endothelial | 0.439 | 0 | 0.010 | 0 | $7.8 \times 10^{-16}$ | $9.2 \times 10^{-16}$ |

*Source data are provided as a source data file.

**Supplementary Table 2.** Key hyperparameters of SigXTalk

| Parameter notation | Description | Default value |
|---|---|---|
| $\boldsymbol{D_{h1}}$ | Dimension of the HGNNConv+, layer 1 | 256 |
| $\boldsymbol{D_{h2}}$ | Dimension of the HGNNConv+, layer 2 | 128 |
| $\boldsymbol{D_{l1}}$ | Dimension of the MLP, layer 1 | 64 |
| $\boldsymbol{D_{l2}}$ | Dimension of the MLP, layer 2 | 32 |
| $\mathbf{Lr_0}$ | Initial learning rate of the optimizer | 0.01 |
| $\boldsymbol{\gamma}$ | Gamma value of the Lr scheduler | 0.975 |
| $\mathbf{Ep}$ | Number of epochs | 50 |
| $\mathbf{Bs}$ | The size of each training batch | 64 |

### Supplementary Notes

#### Supplementary Note 1: Hyperparameter tuning and robustness tests

We test SigXTalk's ability in predicting activated regulatory pathways under different hyperparameter settings using the simulated dataset. The dataset is constructed using SERGIO [1], with the same settings to the dataset used to compare SigXTalk with other methods. Again, the simulated regulatory pathways are divided into training samples and test samples, and the performance is measured by the AUROC and AUPRC values.

Specifically, the effect of initial learning rate ($Lr_0$) and the size of each training batch (batch size, $Bs$) are validated. For the initial learning rate, we set it to 8 different levels from $1 \times 10^{-4}$ to 0.1 and test the performance. For the batch size, we set it to 8 different levels from 16 to 256 and test the performance. For each hyperparameter setting, 5 independent tests are performed.

For the robustness, we test SigXTalk's performance under different experimental conditions, including the number of cells, the number of genes, and the crosstalk levels. The number of cells is set to 5 different levels from 750 cells to 3,750 cells. The number of genes is set to 5 different levels from 100 genes to 500 genes. During this, the ratio of signals:SSCs:targets is fixed to 1:4:15. The density of the gene regulatory network is used to measure the crosstalk levels: intuitively, a denser network implies that a gene is easier to be regulated by other genes, which increases the crosstalk between pathways. Specifically, we set different levels of wiring probability from 0.15 (most sparse) to 1(fully connected) reflect the network density. For each experimental condition setting, 5 independent tests are performed.

For testing the effect of random seeds, we generate the simulated dataset beforehand and manually set 10 random seeds for predicting the activated regulatory pathways. During this test, the dataset, training samples and validation samples remain the same.

To test whether SigXTalk remains effective using self-supervised approaches, we calculate the PPR value for each Signal-SSC pair and the correlation of each SSC-Target pair (as in the analysis of real-world datasets) in the simulated datasets and calculate the $S_H$ value for each Signal-SSC-Target pathway. A proportion (from 20% to 70%) of the pathways with the highest $S_H$ values are selected as the positive training samples and are used to construct the prior hypergraph structure.

#### Supplementary Note 2: Processing of real-world datasets

The gut lineage dataset: the dataset is processed by the data providers. The processing contains quality control, normalization, clustering and annotation, and could be viewed at https://github.com/theislab/gut_lineage. The sample with the highest FVR enrichment (named as the Control_7_FVR_only in the dataset) is used for our analysis.

The HNSCC dataset: we use the Seurat software for the quality control [2]. Specifically, cells with $nCount > 25000$, $nFeature\_RNA > 10000$, or $percent.mt > 2$ are filtered out. Genes that are detected in less than 5% of the cells are filtered out. Next, we identify the genes that are differentially expressed for each cell type using the *FindMarkers* function of the Seurat software (with average $log2fc > 0.25$, adjusted $p - value < 0.001$), and these genes are selected as targets. The subclustering is performed using the *FindSubCluster* function of the Seurat software with the resolution set to 0.25 and the algorithm set to the SLM algorithm[3].

The COVID dataset: we select a COVID patient (sample ID: L17cov) and a healthy control (sample ID: C55ctr), who have the same gender, similar age and similar clinical condition. For the COVID sample, cells with $nCount > 4000$ , $nFeature\_RNA > 4000$, $nFeature\_RNA < 200$, or $percent.mt > 4$ are filtered out. For the healthy control sample, cells with $nCount > 4000$ , $nFeature\_RNA > 3000$, $nFeature\_RNA < 200$, or $percent.mt > 4$ are filtered out. Genes that are differentially expressed in the fibroblasts are selected as targets. These genes are identified using the *FindMarkers* function of the Seurat software (with average $log2fc > 0.25$, adjusted $p - value < 0.001$).

The mouse lung dataset: cells with $nCount > 10000$ , $nFeature\_RNA > 3000$ , or $percent.mt > 7.5$ are filtered out. Genes that are differentially expressed in the fibroblasts are selected as targets. These genes are identified using the *FindMarkers* function of the Seurat software (with average $log2fc > 0.25$, adjusted $p - value < 0.001$).

The mouse brain spatial transcriptomics dataset: the raw sequencing data was processed using Space Ranger. The processed count matrix has been processed with a standard Seurat pipeline and was included as a representative tutorial in Seurat. The clustering and annotation are also provided by Seurat, which uses an ‘anchor’-based integration method to provide a probability distribution of each spot. The label with the highest probability is selected for the cluster that the spot belongs to. Differentially expressed genes (DEGs) of the L6 IT cells are identified using the *FindMarkers* function of the Seurat software. DEGs with significance level $p < 0.001$ are selected as targets.

All the cells of the real datasets used in this research have been clustered and annotated by their providers.

#### Supplementary Note 3: Introduction to hypergraphs and the HGNN+ structure

A hypergraph is represented by a pair $\mathcal{H} = (V, \mathcal{E})$, where $V = \{v_p\}$, $p = 1,2,\dots,N_v$ is the set of nodes and $\mathcal{E} = \{e_q = (v_{q1}, v_{q2}, v_{q3}, \dots)\}$, $q = 1,2,\dots,N_e$ is the set of hyperedges. While the nodes in a hypergraph have nothing different with those in a common graph, a hyperedge may join any number of nodes and thus reflects higher-order interactions, i.e., $e_q \subseteq V$ and $|e_q| \geq 2$, where $|e_q|$ is the number of nodes that $e_q$ contains. For a given node $v_i$, another node $v_j$ is called $v_i$'s neighbor (and vice versa, for undirected hypergraphs), if $\exists\, q$ , $(v_i, v_j) \subseteq e_q$ . Generally, one can characterize a hypergraph structure using the incidence matrix $\mathbf{H} = \{h_{pq}\}$ defined as follows:

$$h_{pq} = \begin{cases} 1, & if\ v_p \in e_q, \\ 0, & if\ v_p \notin e_q. \end{cases} \tag{S1}$$

For a node $v_p$, its node degree is defined as $d(p) \coloneqq \sum_q h_{pq}$, which is quite similar to the node degree in graphs. For a hyperedge $e_q$, its hyperedge degree is defined as the number of nodes it contains, i.e., $\delta(q) \coloneqq \sum_p h_{pq}$. The diagonal matrices of node and hyperedge degrees could then be respectively defined as $\mathbf{D}_v = diag(\{d(p)\})$ of size $N_v \times N_v$ and $\mathbf{D}_e = diag(\{\delta(q)\})$ of size $N_e \times N_e$.

The HGNNConv+ layer used in SigXTalk is realized through the spatial convolution on hypergraph [4]. For a node on the hypergraph, spatial-based hypergraph convolution aggregates its neighbors to update its representations, during which the message is passed from neighbors to the node via the hyperedges targeting it. Formally, we define the node inter-neighbor set $N_v(e)$ as all nodes connected by the hyperedge $e$. Analogously, the hyperedge inter-neighbor set $N_e(v)$ refers to all the hyperedges containing the node $v$. For the $k$-th HGNNConv+ layer, the node features are updated via two steps:

(1) update of the hyperedge features, which could be modelled as follows:

$$\begin{cases} m_e^k = \sum\limits_{v \in N_v(e)} \dfrac{z_v^k}{|N_v(e)|}, \\ y_e^k = w_e \times m_e^k, \end{cases} \tag{S2}$$

where $m_e^k$ denotes the message that the hyperedge $e$ receives, $z_k^v$ denotes the feature of the node $v$, $w_e$ denotes the weight of the hyperedge. In SigXTalk, we simply set $w_e = 1$ so that $y_e^k = m_e^k$. Using the definition of $\mathbf{D}_v$ and $\mathbf{D}_e$, this update could be written in the matrix form:

$$\mathbf{Y}^k = \mathbf{D}_e^{-1}\mathbf{H}^\top\mathbf{Z}^k. \quad (S3)$$

(2) update of the node features, which could be modelled as follows:

$$\begin{cases} m_v^{k+1} = \sum_{e\in N_e(v)} \frac{y_e^k}{|N_e(v)|}, \\ z_v^{k+1} = \sigma\left(m_v^{k+1}\mathbf{\Theta}^k\right), \end{cases} \quad (S4)$$

where $m_v^{k+1}$ is the message that the node $v$ receives, and $x_v^{k+1}$ is the updated node feature of $v$. $\sigma$ represents an activation function like ReLU. $\mathbf{\Theta}_k \in \mathbb{R}^{C^k\times C^{k+1}}$ is the trainable parameter matrix of the $k$-th layer, where $C^k$ denotes the size of feature $z_k^v$. Similarly, we have the following matrix form for this step:

$$\mathbf{Z}^{k+1} = \sigma\left(\mathbf{D}_v^{-1}\mathbf{H}\mathbf{Y}^k\mathbf{\Theta}^k\right). \quad (S5)$$

In all, the update process of the $k$-th HGNNConv+ layer could be written as:

$$\mathbf{Z}^{k+1} = \sigma\left(\mathbf{D}_v^{-1}\mathbf{H}\mathbf{D}_e\mathbf{H}^\top\mathbf{Z}^k\mathbf{\Theta}^k\right). \quad (S6)$$

#### Supplementary Note 4: Implementation of the hypergraph neural network

The *dhg* python package is employed to construct the hypergraph and train the hypergraph neural network [4, 5]. In all, the neural network contains two HGNNConv+ layers, followed by an MLP that consists of two linear layers. Inspired by GENElink [6], the lower-dimensional representation of signal, SSC and target nodes are separately obtained using 3 different MLPs.

While training the neural network, an Adaptive Moment Estimation (Adam) optimizer is used for the stochastic optimization of training loss [7]. As the training proceeds over epochs, a lower learning rate ($\mathrm{Lr}$) is needed to facilitate the convergence of the training loss. Here, an exponential scheduler is adopted for this task. Specifically, the learning rate for the $n$-th epoch could be written as:

$$\mathrm{Lr}_n = \gamma^{n-1}\mathrm{Lr}_0. \quad (S7)$$

Thus, the decay of the learning rate is controlled by the parameter $\gamma$, which is set to 0.975 by default in SigXTalk.

The hyperparameters used during the training process are listed in the Supplementary Table 2.
